# A statistical test for clonal exclusivity in tumour evolution

**DOI:** 10.1101/2021.05.05.442732

**Authors:** Jack Kuipers, Ariane L. Moore, Katharina Jahn, Peter Schraml, Feng Wang, Kiyomi Morita, P. Andrew Futreal, Koichi Takahashi, Christian Beisel, Holger Moch, Niko Beerenwinkel

**Author notes:** Contributed equally (Jack Kuipers, Ariane L. Moore).

## Abstract

Tumour progression is an evolutionary process in which different clones evolve over time, leading to intra-tumour heterogeneity. Interactions between clones can affect tumour evolution and hence disease progression and treatment outcome. Pairs of mutations that are overrepresented in a clonally exclusive fashion over a cohort of patient samples may be suggestive of a synergistic effect between the different clones carrying these mutations. We therefore developed a novel statistical test, called GeneAccord, to identify such gene pairs that are altered in distinct subclones of the same tumour. We analysed our test for calibration and power. By comparing its performance to baseline methods, we demonstrate that to control type I errors, it is essential to account for the evolutionary dependencies among clones. In applying GeneAccord to the single-cell sequencing of a cohort of 123 acute myeloid leukaemia patients, we find 6 clonally exclusive and 2 clonally co-occurring gene pairs. The clonally exclusive pairs mostly involve genes of the key signalling pathways.

## Introduction

Intra-tumour heterogeneity refers to a diverse set of genetically or phenotypically distinct cell populations that coexist within a tumour (Marusyk and Polyak, 2010; Burrell *et al*., 2013). It is the result of mutation, selection, and possibly other evolutionary forces during tumour evolution (Yates and Campbell, 2012; Greaves and Maley, 2012; Beerenwinkel *et al*., 2016). More diverse tumours support more evolutionary pathways and tend to adapt better to treatment and immune responses. Hence they are more likely to escape these selective pressures and develop therapy resistance or immune escape (McGranahan and Swanton, 2015; Dagogo-Jack and Shaw, 2018). Profiling tumours (Irmisch *et al*., 2021) and their heterogeneity therefore has the potential to improve cancer diagnostics and treatment, and the ability to resolve the clonal and subclonal structure of tumours has progressed rapidly with multi-region bulk sequencing (Turajlic *et al*., 2018) and single-cell sequencing (Wang and Navin, 2015; Kuipers *et al*., 2017). Recently, for acute myeloid leukeamia (AML), high-throughput single-cell panel sequencing has uncovered the clonal diversity across two cohorts of 123 patients (Morita *et al*., 2020; Miles *et al*., 2020). These studies show that AML samples tend to have a smaller number of clones, and importantly that multiple different mutations in signalling pathway genes often occur in distinct subclones.

Beyond offering more potential evolutionary pathways, intra-tumour heterogeneity provides more than just a bet-hedging strategy since the individual clones may interact with each other to confer communal advantages (Heppner, 1984; Bonavia *et al*., 2011; Tabassum and Polyak, 2015). Interaction processes between clones are distinct from genetic interactions (epistasis), where, for instance, two genes are mutated in the same cell or clone and together lead to an unanticipated change in the phenotype (Wang *et al*., 2014). The different types of ecological interactions among cancer clones (Tabassum and Polyak, 2015) include negative interactions, for example when clones compete for nutrients or oxygen, while positive interactions such as commensalism, synergism, and mutualism drive tumour cell proliferation and ultimately favour greater intra-tumour heterogeneity. One example of commensalism, where one clone benefits from another one without providing anything in return, is a clone stimulating blood vessel growth, which also supplies other surrounding cells with nutrients and oxygen (Axelrod *et al*., 2006). *Ras*-mutated dermal fibroblast cells have been observed to secrete factors that lead to downregulation of a strong angiogenesis inhibitor in normal cells over 10mm away (Kalas *et al*., 2005), also suggesting that interactions may not only occur between directly neighbouring clones in solid tumours. Cooperation, including synergism and mutualism, where for example, different clones cross-feed each other resources, was hypothesised to be a driving factor in tumour progression (Axelrod *et al*., 2006) and a wealth of examples of clonal cooperation have been discovered in cancer (Miller *et al*., 1991; de Jong *et al*., 1998; Calbo *et al*., 2011; Marusyk *et al*., 2014; Cleary *et al*., 2014; Mateo *et al*., 2014; Chapman *et al*., 2014; Konen *et al*., 2017). If a combination of two or more clones is beneficial, these clones will likely co-exist stably over time and not outcompete each other (Axelrod *et al*., 2006; Cleary *et al*., 2014). More formally, cooperation between cancer clones can be modelled and interpreted using evolutionary game theory (Archetti, 2013; Archetti and Pienta, 2019; Li and Thirumalai, 2020).

As cooperation can play a significant evolutionary role in tumour progression, it is important to elucidate its underlying mechanisms. In particular, understanding how to disrupt this process potentially opens up novel treatment strategies to improve personal cancer treatment. In order to investigate how clones co-exist and possibly interact, a systematic screening of subclonal mutation compositions is necessary. From bulk sequencing data, mutual exclusivity and co-occurrence patterns can be detected from the clonal mutation patterns across cohorts of patients (Jerby-Arnon *et al*., 2014; Kim *et al*., 2015; Leiserson *et al*., 2015; Babur et al., 2015; Constantinescu *et al*., 2016; Cristea *et al*., 2017; Kuipers et al., 2018). This patient-level resolution however does not account for the subclonal structure within each patient overlooking potential subclonal interactions.

Resolving tumours at the subclonal level with multi-region bulk or single-cell sequencing offers a route to address this. However it involves another challenge: the genotypes of the subclones are not independent observations. Hence, treating clones like tumour samples in the above-mentioned patient-level analyses would lead to spurious correlations. Instead, we must account for the dependency structure among clones encoded by their phylogenetic relationships. For example, for two mutations in a common lineage, after the first mutation we have clones with the genotype (1, 0) resembling an exclusivity pattern while after both mutations we have the genotype (1, 1) indicating co-occurrence. To avoid treating the dependency structure, previous analyses have, for example, been limited to the common ancestor clone of the mutations to ensure independence, as was the case for the analysis of clonal co-occurrences and exclusivities of ten selected driver events in clear cell renal carcinoma using multi-region bulk sequencing (Turajlic *et al*., 2018). Here we develop a statistical test, called GeneAccord, which considers the full phylogenetic tree representing the complete evolutionary history of each tumour and the subclonal mutation patterns across the entire tree. GeneAccord can additionally take into account the uncertainty in the tree inference by allowing the input of multiple alternative tree topologies per patient. With this input, GeneAccord assesses, in a statistically rigorous way, whether specific sub-clonal mutation combinations occur at a higher or lower rate than expected by chance over the cohort of patient tumour samples. We evaluate GeneAccord’s power and demonstrate that it correctly controls the type I error rate, unlike baseline alternatives which do not account for the underlying phylogenetic trees. Finally, we illustrate GeneAccord on a cohort of 123 AML patients resolved with single-cell sequencing (Morita *et al*., 2020). We find significant signs of clonal exclusivity, particularly between genes involved in signalling pathways.

## Results

### GeneAccord algorithm

In order to systematically analyse subclonal mutation combinations in tumour clones, we developed a statistical framework and implemented it in the R package GeneAccord. The statistical test can identify pairs of mutated genes or pathways that both occur in the same tumour but in different clonal lineages. We refer to the situation of one clone lineage possessing mutations that the other one does not have, and vice versa, as clonal exclusivity. By resolving the evolutionary history of tumours, for example through single-cell sequencing and reconstructing the genotypes of the clones, clonally exclusive gene pairs will display mutual exclusivity (Figure 1). The underlying rationale for searching for clonally exclusive gene pairs is that if two clones co-exist in a tumour and cooperate, for example, by sharing diffusible factors, they may have acquired complementary sets of mutations for mutual benefit.

**Figure 1:**
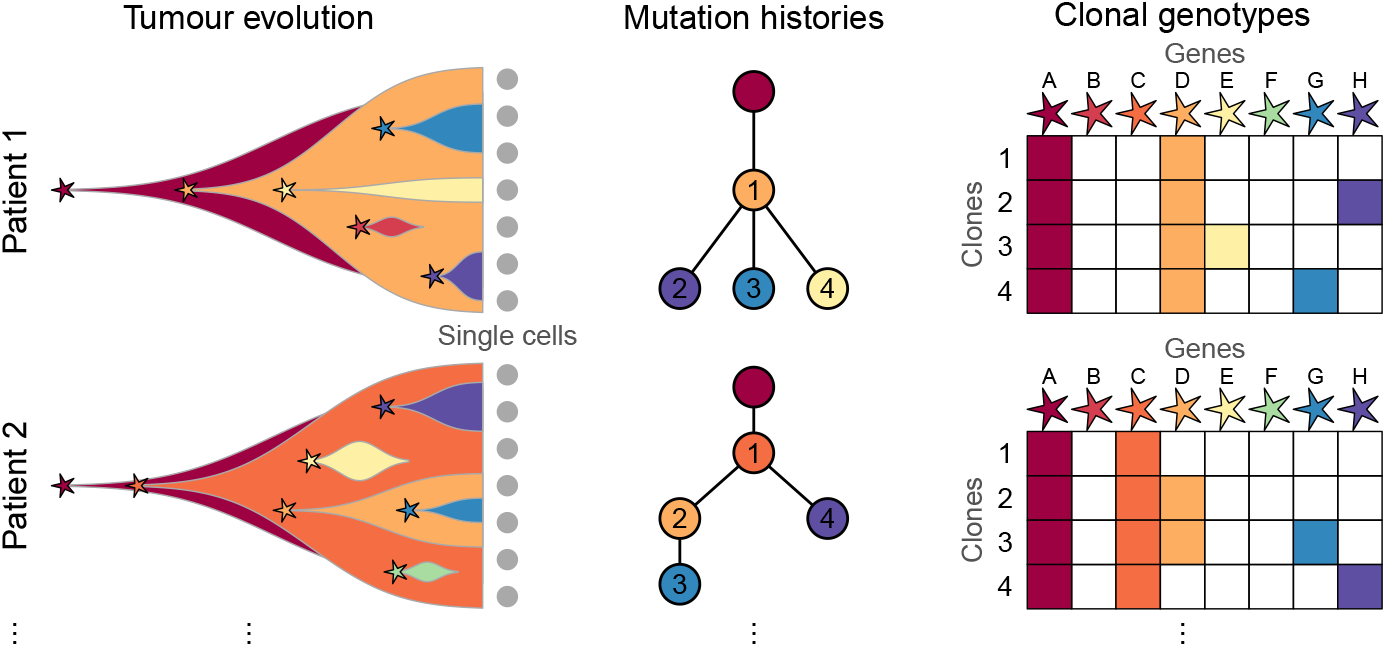
Tumour evolution and clonal exclusivity. During tumour evolution different mutations may arise leading to heterogeneous subclones with distinct genotypes. From single-cell sequencing we may reconstruct the mutational history of each tumour, encoding the ordering of mutations and their phylogenetic relationships. The clones are numbered in each tree and inherit mutations in ancestral clones. For example in patient 1, clone 2 exhibits mutations in genes A, D and H (red, orange, purple) as displayed in the clone-genotype matrix on the right. Clonally exclusive mutations will appear in different branches of the trees and exhibit mutually exclusive patterns in the clone-genotype matrices, as exhibited for example for the two rightmost mutations G and H (blue and purple) in both patients.

To search for clonal exclusivity, we evaluate across a cohort of patient tumours whether the observed patterns of exclusivity occur more or less often than expected by chance alone. Specifically, we compare to the null model of mutations being placed randomly within each tumour phylogeny. We frame the testing procedure as a likelihood ratio test with a single clonal exclusivity score Δ for each gene pair, which when negative indicates that the gene pair is mutated in different clones more often than expected, and when positive indicates a higher rate of co-occurrence in the same clonal lineage. We developed and implemented the statistical test as an exact and asymptotic test in the GeneAccord R package (Methods, https://github.com/cbg-ethz/GeneAccord). After testing gene pairs, multiple testing correction is performed to control the false discovery rate with the Benjamini-Hochberg procedure (Benjamini and Hochberg, 1995). If a pair is significantly clonally exclusive, it suggests that this specific clone configuration may confer a selective advantage, possibly through cooperation between the clones.

### Calibration and power of the test

The statistical testing (Methods) is a likelihood ratio test where the test statistic asymptotically follows a chi-squared distribution. For gene pairs observed in a smaller number *n* of patient samples this asymptotic result can however be poorly calibrated. For example, we simulated data under the null where the background clonal exclusivity rates *r* are sampled from a beta distribution with parameters 2 and 3 which gives a similar mean and variance to the rates computed from the real AML dataset. Using the chi-squared distribution to compute p-values under the null we see an enrichment of lower p-values for smaller *n* (Figure 2). The chi-squared approximation however seems appropriate when the gene pair is observed in enough samples (more than ~10).

**Figure 2:**
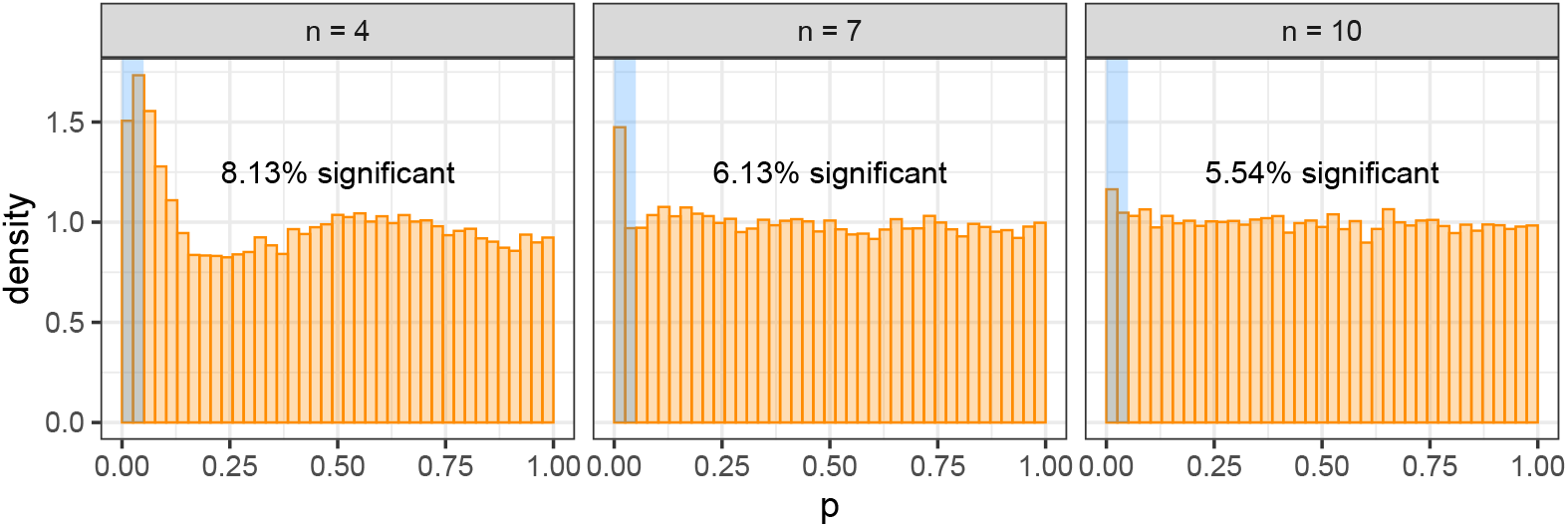
Calibration of the chi-squared approximation. For gene pairs observed in a smaller number *n* of patient samples, we observe overly liberal p-values and a lack of calibration. For larger *n*, the approximation becomes more appropriate.

We therefore developed an exact version of the test (Methods) to ensure calibration. For the same simulated example as before we now observe fewer than 5% of p-values under the null being significant at a 5% threshold (Figure 3). However, since the number of possible outcomes is limited, we also observe very strong discrete effects in the null p-value distribution.

**Figure 3:**
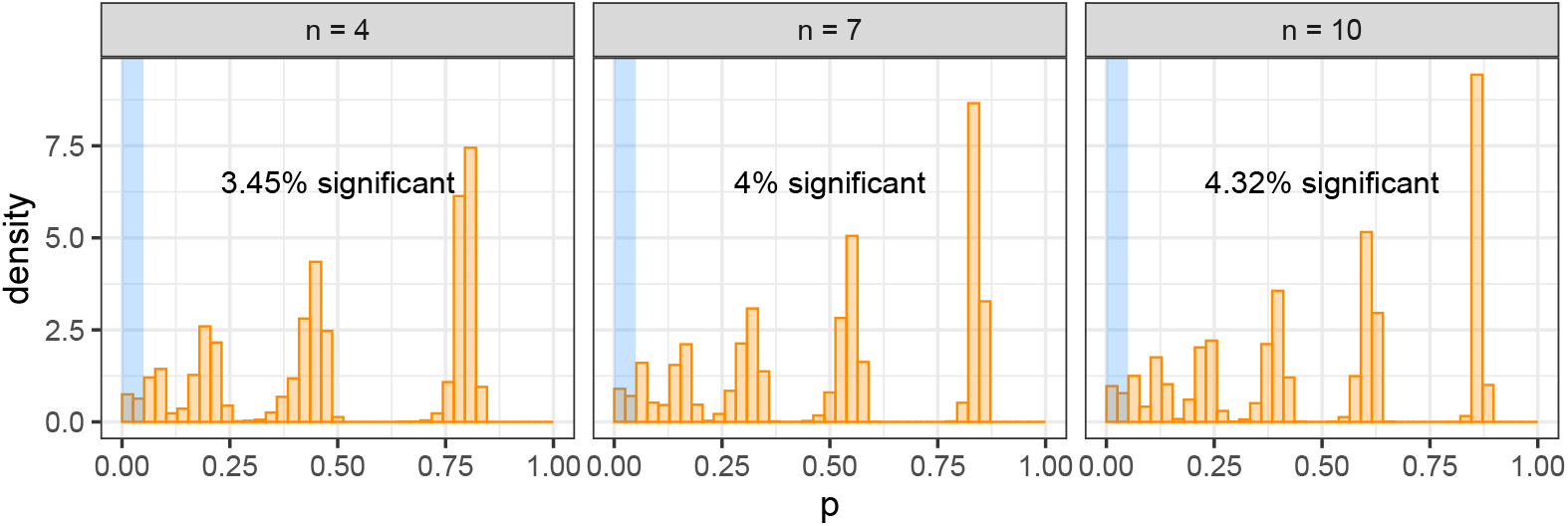
Calibration of the exact test. For the exact test we observe conservative p-values at lower significance levels, and very pronounced discrete effects for larger values.

For larger *n*, enumerating all possible outcomes for the exact test becomes more computationally expensive, while the chi-squared approximation improves. As a default in the R package we therefore switch to the chi-squared approximation for *n* > 12.

To check calibration while dampening the discrete effects, we created a Monte Carlo version of the exact test (Methods) into which we could additionally add smoothing through adding noise to the sampled clonal exclusivity rates. Performing such smoothing (with *ν* = 10) we observe uniformity of the p-values under the null for larger *n*, with some remnants of the discrete effects visible for lower *n* (Figure 4). The simulated data therefore show that the exact test is calibrated on average.

**Figure 4:**
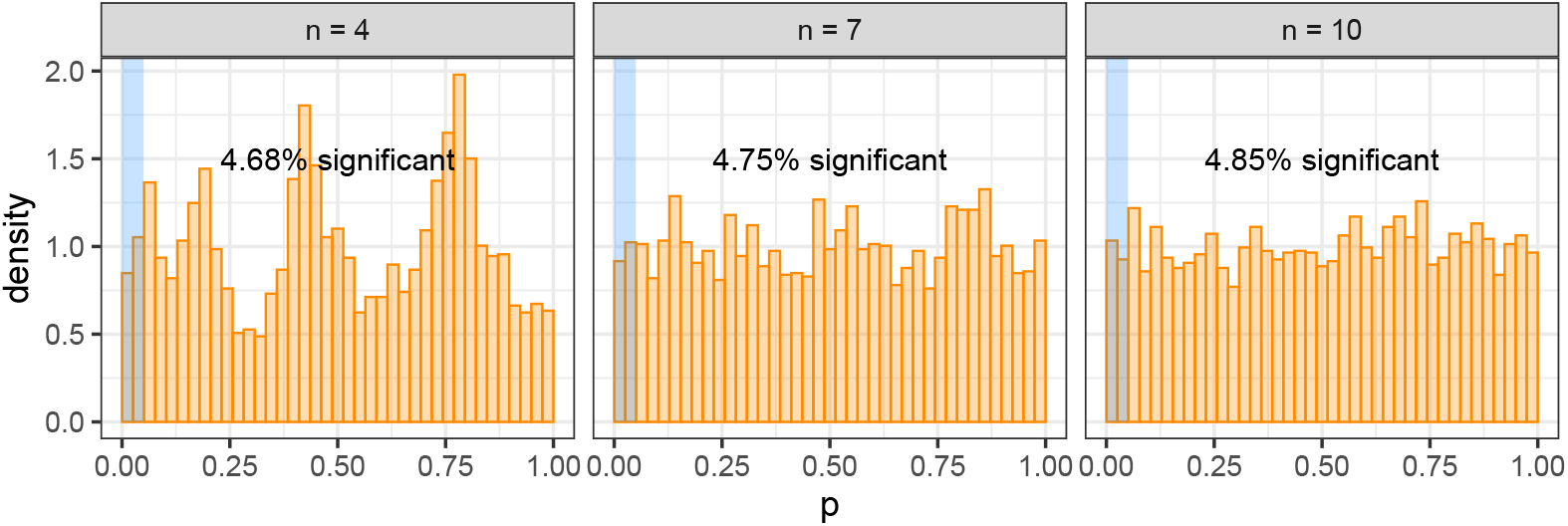
Calibration of the exact test with Monte Carlo smoothing. Smoothing the exact test by adding noise to the rates, we observe good calibration and the desired uniform distribution of p-values under the null for the larger *n*.

Having checked the calibration of the GeneAccord exact test under the null, we next evaluate its power by simulating data under the alternative for different values of Δ. The rates are again sampled from a beta distribution with parameters 2 and 3. For very small sample sizes, we need a relatively strong effect to have a high power (for example a Δ of −4 to have a power of 75% for gene pairs in only 4 patients), though the power rapidly increases for larger samples sizes as expected; for example, a Δ of −3 can be detected in 10 patients with a probability of over 96% (Figure 5).

**Figure 5:**
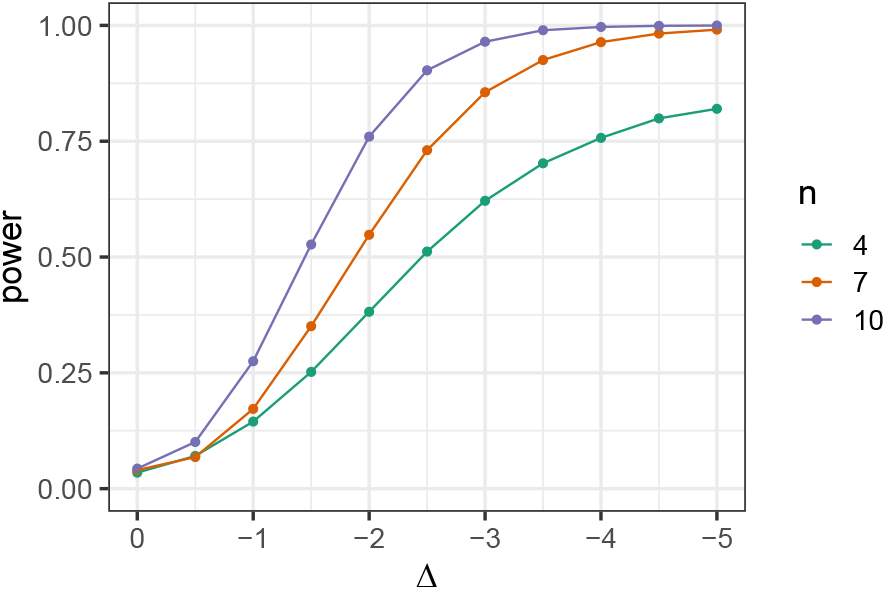
Power of the exact test. The power of the exact test as we increase the effect size for different sample sizes.

### Comparison to exclusivity testing without trees

Alternative methods to GeneAccord to test for exclusivity do not take the phylogenetic information into account. As a demonstration that utilising evolutionary histories is necessary, we perform naïve exclusivity testing at the clonal level by constructing a contingency table for each gene pair and running standard independence testing: the Fisher’s exact test, the G-test using the chi-squared distribution and the log odds ratio test with the normal approximation. To generate data we sampled random binary trees each with 10 inner branches and uniformly distributed 20 mutations across those branches to create 10 clonal genotypes per tree. We collated sets of 10 trees, corresponding to gene pairs observed in 10 patient samples, and ran the standard independence tests along with GeneAccord. The entire procedure was repeated 400 times.

As the mutations are interchangeable in their random placement in the trees, we are in the null setting of no clonal enrichment. The standard tests, however, are very heavily confounded and find significant results (at the 5% level) roughly half the time (Figure 6, top row). Not taking the underlying tree structure into account, and treating the clonal genotypes as independent observations, therefore leads to spurious exclusivity and co-occurrence patterns. GeneAccord’s exact test remains properly calibrated, as evidenced with the Monte Carlo smoothing, while there is a slight enrichment with the chi-squared approximation (Figure 6, bottom row).

**Figure 6:**
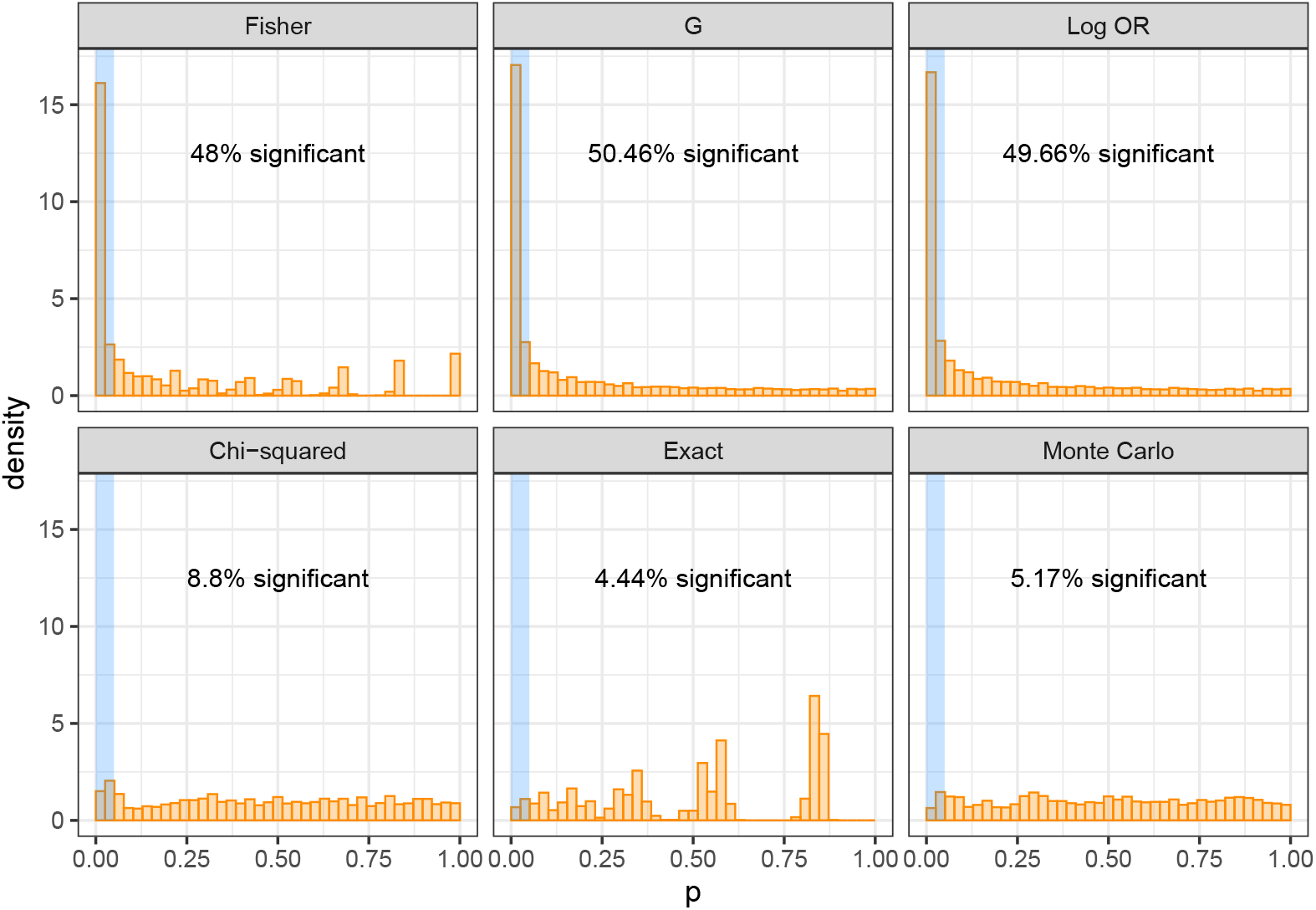
Comparison to standard exclusivity tests. Standard independence tests (top row: Fisher’s exact test, G-test and log odds ratio (OR) test, from left to right) are heavily miscalibrated for data generated from placing mutations on lineage trees under the null. GeneAccord (bottom row: chi-squared approximation, exact test and with Monte Carlo smoothing, from left to right), apart from some enrichment with the chi-squared approximation, shows proper calibration.

### AML gene pairs

We ran the GeneAccord algorithm on the cohort of 123 AML patient samples from Morita *et al*. (2020) whose evolutionary histories have been reconstructed with the single-cell phylogeny method SCITE (Jahn *et al*., 2016). The results, summarised in Table 1, show 6 gene pairs with significant evidence of clonal exclusivity, and 2 gene pairs with co-occurrence.

**Table 1:** GeneAccord results for the AML cohort (Morita *et al*., 2020). Ranked list of the gene pairs tested with the GeneAccord algorithm on the cohort of 123 AML patient samples. For each gene pair, the column *n* is the number of patients exhibiting both gene mutations, *n*_cx_ the number of times the genes are clonally exclusive. Δ is the clonal exclusivity score indicating enrichment of clonal exclusivity (negative) or clonal co-occurrence (positive). LLR is the log-likelihood ratio statistic, p is the p-value and q the adjusted p-value after Benjamini-Hochberg correction. Linear trees which are not informative for the testing have been excluded. Only gene pairs in more than 3 patient samples are included.

| Rank | Gene pair | $n$ | $n_{cx}$ | $\Delta$ | LLR | $p$ | $q$ |
| --- | --- | --- | --- | --- | --- | --- | --- |
| 1 | *NRAS_PTPN11* | 8 | 8 | $-\infty$ | 141.64 | 0.00042 | 0.0088 |
| 2 | *FLT3_NRAS* | 8 | 7 | -2.704 | 93.78 | 0.0021 | 0.022 |
| 3 | *NPM1_PTPN11* | 8 | 0 | ∞ | 90.40 | 0.0059 | 0.034 |
| 4 | *KRAS_NRAS* | 9 | 8 | -2.434 | 77.00 | 0.0069 | 0.034 |
| 5 | *KRAS_PTPN11* | 4 | 4 | $-\infty$ | 79.94 | 0.0092 | 0.034 |
| 6 | *FLT3_KRAS* | 4 | 4 | $-\infty$ | 76.86 | 0.011 | 0.034 |
| 7 | *FLT3_NPM1* | 11 | 0 | ∞ | 81.31 | 0.011 | 0.034 |
| 8 | *IDH1_IDH2* | 4 | 4 | $-\infty$ | 67.46 | 0.017 | 0.045 |
| 9 | *KRAS_NPM1* | 5 | 0 | ∞ | 58.62 | 0.040 | 0.092 |
| 10 | *FLT3_PTPN11* | 7 | 5 | -1.756 | 41.64 | 0.055 | 0.115 |
| 11 | *NPM1_NRAS* | 11 | 2 | 1.321 | 29.17 | 0.129 | 0.246 |
| 12 | *IDH2_PTPN11* | 4 | 0 | ∞ | 31.22 | 0.191 | 0.303 |
| 13 | *PTPN11_WT1* | 5 | 3 | -1.469 | 20.28 | 0.193 | 0.303 |
| 14 | *DNMT3A_FLT3* | 4 | 0 | ∞ | 24.64 | 0.202 | 0.303 |
| 15 | *IDH2_NRAS* | 6 | 3 | -0.828 | 0.908 | 0.270 | 0.379 |
| 16 | *IDH2_NPM1* | 7 | 1 | 1.098 | 12.47 | 0.314 | 0.413 |
| 17 | *DNMT3A_NPM1* | 5 | 1 | 0.618 | 0.326 | 0.495 | 0.578 |
| 18 | *IDH1_NPM1* | 5 | 1 | 0.847 | 0.635 | 0.496 | 0.578 |
| 19 | *DNMT3A_IDH2* | 4 | 1 | 0.521 | 0.179 | 0.780 | 0.804 |
| 20 | *FLT3_IDH1* | 4 | 1 | 0.297 | 0.066 | 0.791 | 0.804 |
| 21 | *FLT3_IDH2* | 5 | 1 | 0.407 | 0.137 | 0.804 | 0.804 |

One clonally exclusive gene pair is between the two *IDH* genes, while the other 5 involve the genes *FLT3, NRAS, KRAS* and *PTPN11*, which affect the receptor tyrosine kinase (RTK)/Ras GTPase (RAS)/MAP Kinase (MAPK) signalling pathways. Clonal exclusivity would align with functional redundancy making the mutations interchangeable, though having several mutations in parallel lineages is evolutionarily more complex than sharing a single mutation in an ancestral clone. The significant clonal exclusivity of these gene pairs may then point to stronger effects like cooperation across the clones or synthetic lethality where having co-occurring mutations in the same lineage leads to strong decrease in viability of the tumour clone (Unni *et al*., 2015; Varmus *et al*., 2016).

We observe significant co-occurrence of *NPM1*, the most frequent mutation in the cohort, with both *PTPN11* and *FLT3*. There is a strong interplay between *NPM1, FLT3* and age in terms of survival prognosis (Juliusson *et al*., 2020), and proven biological cooperation in AML between the genes (Mupo *et al*., 2013; Mallardo *et al*., 2013; Rau *et al*., 2014).

## Discussion

We have introduced GeneAccord as a novel statistical framework to systematically analyse the subclonal mutation patterns in cohorts of tumour patients. By introducing a null model of the mutations randomly placed in the evolutionary histories of each patient, we created a new exact test to assess how unusual the subclonal patterns are across the cohort. We evaluated the calibration and power of the test. For larger numbers of patient samples exhibiting the same pair of genes, the exact test becomes more computationally intensive, while utilising the asymptotic chi-squared distribution becomes better calibrated and is computationally cheap. We focussed on clonal exclusivity at the gene level, but the method applies to any marker that can be assigned to the clones in the architecture of each tumour, for example to pathways by mapping from mutations to pathways (Moore *et al*., 2021) or from differentially expressed genes to pathways.

We detect pairs of genes or pathways that have an elevated or depressed rate of mutating in different clonal lineages of the same tumour. There are several possible biological explanations for such mutational patterns. For clonally exclusive gene pairs occurring in separate co-existing clones, the two clones could have complementing phenotypes and cooperate or mutually benefit each other, for instance, by sharing diffusible factors. Another possibility is that the two genes of a clonally exclusive pair are synthetically lethal and that both genes being mutated in the same cell would lead to a disadvantageous, and maybe even lethal, phenotype. The two clones may also be the result of parallel convergent evolution where both clones exhibit the same phenotype by different mutations. The specific subclonal mutation pattern may also depend on the evolutionary subtype (Turajlic *et al*., 2018). In order to gain a better understanding of possible reasons for the clonally exclusive pattern, it is important to examine the biological functions of these genes and pathways in more detail. A definite proof of such interactions then requires future experimental studies in vitro or in vivo. Here, we have developed and applied a new computational approach to identify such gene (or pathway) pairs that are unlikely to be generated by random chance alone. As such, the GeneAccord method will be useful in finding and prioritising candidate cooperative tumour clones in cancer patient cohorts.

While many cancer-related mutually exclusive gene pairs have been previously identified across cohorts at a bulk or consensus level, finding such pairs within a single tumour and its clonal architecture has received less attention. Especially with the high resolution of multi-region bulk and single-cell sequencing we can now better reconstruct the evolution histories of tumours for downstream analyses as performed here. As the panels used for high-throughput single-cell sequencing increase in size and coverage we can expect more detailed tumour phylogenies and better clonal resolution to provide more power to detect clonal exclusivity patterns. For example, in the current cohort of AML patients many exhibit only a handful of mutations among the panel sequenced and often in a linear order. These samples are not informative for our exclusivity testing reducing our effective sample size. On a related front, our testing is conditioned on the observed tumour phylogenies, but the phylogenetic structure itself may be the result of clonal interactions and shed further light on them. As a future extension we could integrate our testing with evolutionary modelling to extract additional signals of subclonal cooperation. In particular, linear trees are uninformative for the GeneAccord test, whereas such topologies may be favoured with co-occurring gene pairs. Currently we focused on testing genes pairwise, and another possible extension would be to consider larger sets of genes and detect more generalised (higher order) clonal exclusivity patterns.

Looking ahead, combined single-cell sequencing of the exome and transcriptome (Macaulay *et al*., 2015) will allow the assignment of differentially expressed genes to specific clones. Alternatively, matching multi-omic profiling of the different cells from the same sample, for example, through matching expression profiles and copy number states (Campbell *et al*., 2019) on evolutionary trees (Kuipers *et al*., 2020), would provide a transcriptomic profile for each clone. This would enable a GeneAccord analysis on the transcriptomic level. Therefore, with single-cell data, one could perform a combined analysis using various omics layers including differentially expressed genes, as well as copy number and epigenetic changes. This would allow for a more holistic portrait of the subclonal alteration profiles and potentially reveal more synergies between clones that could inform the design of future treatments.

## Methods

### GeneAccord overview

The input data to the GeneAccord algorithm are the mutated gene-to-clone assignments from a cohort of cancer patients. These are obtained by running phylogenetic tree inference methods, for example on single-cell sequencing data. We utilised the trees inferred by SCITE (Jahn *et al*., 2016) for the cohort of 123 AML patient samples (Morita *et al*., 2020).

In general, there is uncertainty in the tree structure learned from sequencing data. Therefore, our algorithm was designed to allow as input multiple gene-to-clone assignments per patient, for example by sampling from the posterior distribution of trees. The tree inference designates point mutations to individual clones, while mutations can then be mapped to genes or pathways using existing pathway databases. Our statistical framework can be applied on the gene level, or on the pathway level to detect clonally exclusive pairs of pathways. We focus on the gene case here.

### Likelihood ratio test

To test if gene pairs have a different rate of clonal exclusivity compared to typical genes, we first compute the background rate of clonal exclusivity for each patient *i*

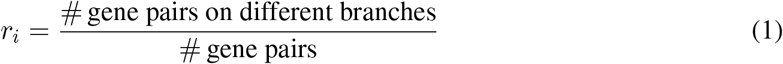

When we have a sample of trees for the patient, we average this rate across the trees. Then for each gene pair (*j*, *k*), we look at the clonal exclusivity of gene *j* and gene *k* among the patient samples that possess both somewhere in their trees. For each patient *i*, we compute

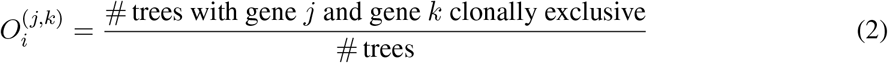

When we have a single tree for patient *i*, this quantity will be either 0 or 1. Under the null model that genes are placed randomly on the trees for each patient, the likelihood that a gene pair will be exclusive is *r_i_* while the likelihood of co-occurrence is (1 – *r_i_*). The log-likelihood of the observed exclusivity patterns of the gene pair is then

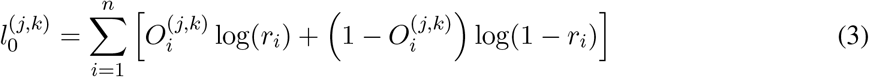

where the sum is only over patients with both genes present. For the alternative model, we allow the clonal exclusivity rates to differ for the gene pair. To ensure that the shifted rates are between 0 and 1, we perform the shift in the logit space and define the shifted rate 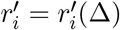 by

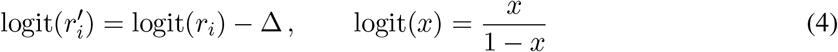

where the sign of the shift Δ is chosen so that negative values indicate clonal exclusivity and positive values indicate co-occurrence. Then we maximise the log-likelihood of the alternative over Δ

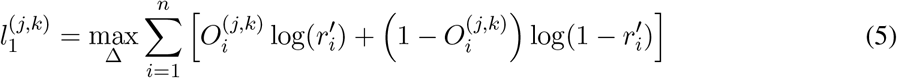

where the dependence on Δ is through the *r*′ and the maximisation is performed numerically (with the optimize function in R). For computational reasons, we restrict Δ to a range of ±10 since after the logit transformations the *r*′ will essentially be 0 or 1 and further increasing or decreasing Δ to ±∞ will hardly affect the maximal log-likelihood numerically. As a test statistic we employ the log-likelihood ratio (LLR)

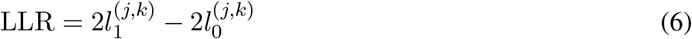

For linear trees all gene pairs are in the same lineage and with none being clonally exclusive *r_i_* = 0. The logit transform maps to −∞ which is unaffected by the shift of Δ and the transformed rate 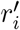 will also be 0. Similarly, for star trees with every gene in its own lineage and *r_i_* = 1, the shifted rates are unchanged. These topologies therefore do not contribute to the LLR statistic, and patients with such uninformative trees are removed from the cohort in a pre-processing step. If we denote by *n* the number of patients samples with both genes, for larger *n* the LLR statistics will follow a 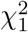 distribution.

### Exact test

To be able to use GeneAccord for less common gene pairs we devise an exact test for smaller *n*. When the gene pair is observed in *n* patient samples, there are 2^*n*^ different possible binary clonal exclusivity patterns. If we store the binary pattern in a vector *b*, the probability of it arising under the null is

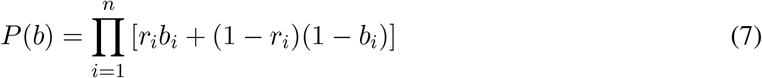

For each binary vector, we compute its LLR statistic. The p-value is the sum of the probabilities of the binary vectors with a LLR statistic larger than the observed statistic. In the p-value we also include half the probabilities of binary vectors with identical LLR statistics. This is necessary to obtain calibration of the p-values, which we demonstrate with Monte Carlo smoothing.

### Monte Carlo smoothing

To better check the calibration of the exact test we wish to introduce some smoothing into the p-value distribution. Rather than enumerating all binary vectors, we could simply sample them proportionally to their probabilities and obtain a Monte Carlo estimate of the p-value. In the limit of an infinite sample size this reduces to the value from the exact test, with the same discrete effects. To smooth these effects we add some noise to the values of *r* for each Monte Carlo sample by sampling them from a beta distribution with parameters *νr* and *ν* (1 – *r*). The parameter *ν* corresponds to the amount of overdispersion or noise in the sampled rates with the limit *ν* → ∞ being noiseless. From the Monte Carlo samples, the p-value is the proportion of samples with a larger LLR statistic (the probability of being equal is 0). As we take the limit *ν* → ∞ the median of a beta distribution approaches the mean, so that half the Monte Carlo samples which would have the same LLR in the exact test will be more extreme, and half less extreme. In the exact test we therefore count binary vectors with the same discrete LLR statistic with weight half.

### Computational cost

After the data pre-processing, computing the LLR statistic involves optimising the log-likelihood numerically while each likelihood computation is linear in *n*, the number of patient samples. For the chi-squared approximation we perform a single optimisation so for each gene pair considered the cost is *O*(*n*). The exact test involves optimising 2^*n*^ log-likelihoods leading to a cost of *O*(*n*2^*n*^). The Monte Carlo smoothed version optimises for each Monte Carlo sample, so with *N* samples the cost is *O*(*nN*).

### Alpha budgeting

For gene pairs mutated in only a handful of patient samples, depending on the background clonal exclusivity rates for those patients, it may never be possible to get a significant p-value below 5% regardless of the observed clonal exclusivity patterns in the data. For each gene pair we therefore compute, from the set of patient samples possessing both mutations, the minimum possible p-value. If the minimum is greater than 5%, the gene pair is removed before the testing with GeneAccord so as not to affect multiple testing corrections.

### AML cohort processing

In the AML cohort (Morita *et al*., 2020), we start with the trees inferred with SCITE (Jahn *et al*., 2016) and filtered out any clones with a frequency below 1%. Since most patient samples only exhibit a few mutations amongst the panel of 30 genes selected, the majority of trees are linear. Of the 123 patient samples, 35 have trees which are informative and these are retained for the GeneAccord analysis. We then consider gene pairs which were both present in at least 4 patient trees, of which there are 22 pairs. For one pair there was no power to detect significance so it was removed, leaving a final set of 21 gene pairs (Table 1).

## Availability of data and materials

The original data was previously published (Morita *et al*., 2020) and the processed data along with the analysis with GeneAccord are available at https://github.com/cbg-ethz/GeneAccord

## Author contributions

Conceptualization: JK, ALM, CB, PS, HM, NB; Data Curation: KJ, FW, KM, PAF; Formal Analysis: JK, ALM; Methodology: JK, ALM, NB; Resources: PS, FW, KM, PAF, KT, CB, HM; Software: JK, ALM, KJ; Supervision: PS, KT, CB, HM, NB; Visualization: JK, ALM; Writing – Original Draft Preparation: JK, ALM, KJ, KT, NB; Writing – Review & Editing: all authors.

